# Maresin-1 promotes neuroprotection and prevents disease progression in experimental models of multiple sclerosis through metabolic reprogramming and shaping innate and adaptive disease-associated cell types

**DOI:** 10.1101/2023.09.25.559216

**Authors:** Insha Zahoor, Mohammad Nematullah, Mohammad Ejaz Ahmed, Mena Fatma, Sajad Mir, Kamesh Ayasolla, Mirela Cerghet, Suresh Palaniyandi, Veronica Ceci, Giulia Carrera, Fabio Buttari, Diego Centonze, Yang Mao-Draayer, Ramandeep Rattan, Valerio Chiurchiù, Shailendra Giri

## Abstract

Multiple sclerosis (MS) is one of the most common inflammatory neurodegenerative diseases in young adults and causes neurological abnormalities and disability. We studied the effect of maresin 1 (MaR1) on the progression of disease in a relapsing-remitting form of experimental allergic encephalomyelitis (RR-EAE). Treatment with MaR1 in RR-EAE accelerated inflammation resolution, protected against neurological deficits, and delayed disease progression by decreasing immune cell infiltration (CD4+IL17+ and CD4+IFN*γ*+) into the CNS. Furthermore, the administration of MaR1 increased the production of IL-10, predominantly in macrophages and CD4+ cells. However, neutralizing IL-10 with an anti-IL-10 antibody abolished the protective effect of MaR1 on RR-EAE, suggesting that IL-10 plays a role in mediating the protective effect of MaR1 on EAE. Metabolism is rapidly becoming recognized as an important factor influencing the effector function of many immune cells. Using cutting-edge metabolic assays, our study revealed that compared with vehicle treatment, MaR1 treatment effectively restored the metabolic dysregulation observed in CD4+ cells, macrophages, and microglia in the treated group. Furthermore, MaR1 treatment reversed defective efferocytosis in EAE mice, which was potentially facilitated by the induction of metabolic alterations in macrophages and microglia. MaR1 treatment also protected myelin in the EAE group and regulated the metabolism of O4+ oligodendrocytes by restoring metabolic dysregulation through improved mitochondrial function and decreased glycolysis. Overall, in a preclinical MS animal model, MaR1 treatment produced anti-inflammatory and neuroprotective effects. It also triggered metabolic reprogramming in disease-associated cell types, accelerated efferocytosis, and preserved myelination. These data support that MaR1 has potential as a novel treatment agent for MS and other autoimmune diseases.

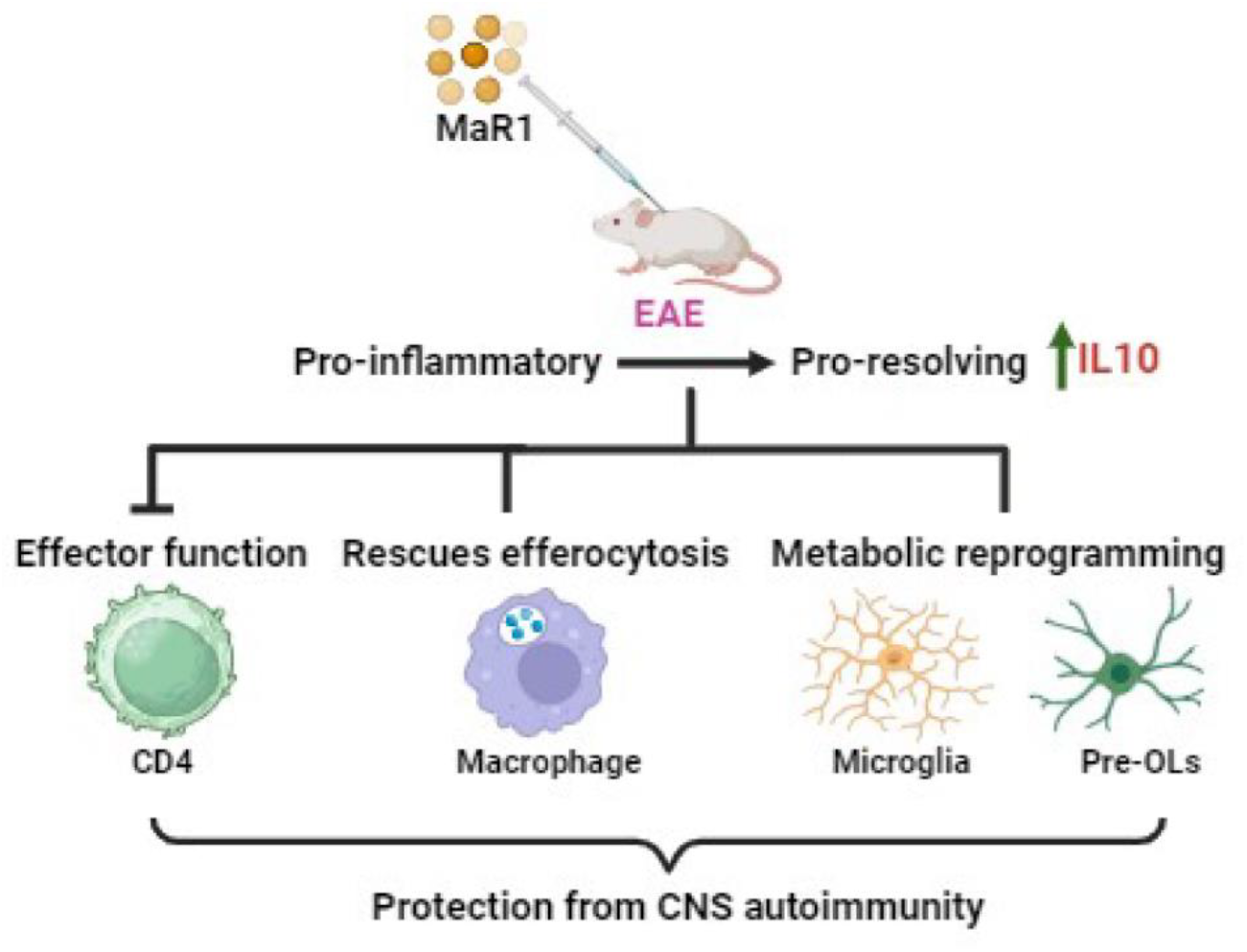

**Highlights:**

- MaR1 expedited inflammation resolution and prevented neurological impairments in RR-EAE.
- IL-10 plays a role in mediating the protective effect of MaR1 on EAE.
- MaR1 repaired CD4, macrophage, and microglia metabolic abnormalities.
- MaR1 therapy restored efferocytosis in EAE.
- MaR1 preserved myelin and improved O4+ oligodendrocyte metabolism.

## Introduction

Multiple sclerosis (MS) is one of the most common inflammatory and neurodegenerative diseases in young adults and leads to the development of neurological defects accompanied by irreversible disability [1, 2]. Although the relapsing-remitting phenotype of MS is treatable, overall, the disease remains largely incurable and results in progressive disability. Patients commonly suffer from relapses accompanied by damage-inducing inflammation, and current immunomodulatory medications suppress the whole immune system, resulting in ill-tolerated side effects and subpar tolerability [3]. Inflammation plays a detrimental role in several autoimmune diseases, including MS, making it important to investigate endogenous inflammatory pathways [4]. During the disease process, the human body is pushed into remission or chronic disease by an imbalance of inflammation and endogenous defense mechanisms. The endogenous mechanisms involved in combatting inflammation in MS patients are not sufficient to resolve this disease, resulting in chronic inflammation. Unresolved inflammation represents the pathological hallmark of MS and several other autoimmune diseases; however, current therapeutic options fail to adequately suppress ongoing inflammation, resulting in inflammatory attacks that gradually increase in severity, causing continued neuronal damage and failed or inadequate remyelination, consequently leaving axons demyelinated and vulnerable to degeneration [5, 6]. Therefore, investigating the pathways that can reduce excessive inflammation and promote rapid inflammation resolution is highly important.

Inflammation resolution is an active process mediated by endogenously derived metabolites of dietary polyunsaturated omega fatty acids (PUFAs), which are collectively termed specialized proresolving lipid mediators (SPMs). Dietary factors are modifiable environmental contributors to MS [7, 8]. A large prospective study revealed a significant inverse association between polyunsaturated fatty acid (PUFA) intake and the risk of MS [9]. Thus, low PUFA intake may be a modifiable risk factor for MS. In-depth research on dietary factors is needed to determine the specific mechanisms of disease etiopathogenesis. There is compelling evidence suggesting defects in docosahexaenoic acid (DHA) metabolism, which results in the deficiency of downstream SPMs (resolvins, protectins, and maresins), leading to chronic inflammation and a delay in the healing/repair process [10, 11]. However, limited information is available regarding the presence or abundance of these metabolites in MS patients [12].

Our group previously demonstrated that not only the biosynthesis of the DHA-derived metabolite 14-HDoHE, a pathway marker for SPM maresin, is significantly lower in patients with RRMS [13] but also that MaR1 levels are undetectable in the plasma of both relapsing-remitting and progressive MS patients [14]. MaR1 is known to mitigate inflammatory damage in other inflammatory disease models, with promising neurological recovery achieved by promoting resolution and neuroprotection and attenuating neuroinflammation and neurocognitive dysfunction [15-17]. As a new family of anti-inflammatory and proresolving lipid mediators, MaRs are a class of 14S-dihydroxyl-containing molecules with conjugated triene double bonds that are synthesized from DHA through an oxidative (e.g., lipoxygenase-related) pathway during inflammation regression. They are highly conserved resolution mediators with potent anti-inflammatory and proresolving properties and play a role in tissue regeneration in acute or chronic inflammatory-related disease models [18-20]. Therefore, MaR1, a potent inflammatory self-limiting factor, is expected to become a highly promising anti-inflammatory intervention drug target. Briefly, MaR1 is synthesized from DHA by 12-LOX, with 14-HpDHA serving as its precursor and 14-HDHA serving as a byproduct (pathway marker). It binds to the receptor leucine-rich repeat containing G protein-coupled receptor 6 (LGR6), and further studies are needed to elucidate whether the 12-LOX/14-HDHA/LGR6 pathway is a crucial metabolic signaling axis involved in the regulation of MaR1 bioaction in peripheral inflammation and neuroinflammation and, if so, may provide us with attractive therapeutic target(s) with potential clinical application for MS treatment.

The current study sought to establish a comprehensive protective mechanism for MaR1 in a relapsing-remitting (RR) animal model of experimental allergic encephalomyelitis (EAE) and in blood obtained from RR-MS patients.

## Materials and Methods

### EAE induction and functional evaluation

Female 10-to 12-week-old SJL mice were purchased from the Jackson Laboratory (Bar Harbor, ME). The animals were housed in the pathogen-free animal facility of Henry Ford Health, Detroit, MI, according to the animal protocols approved by the Animal Care and Use Committee of Henry Ford Hospital. Mice were immunized on day 0 by subcutaneous injection in the flank region with a total of 200 μl of the antigen PLP_139-151_ peptide (200 μg/mouse) emulsified in Complete Freund’s Adjuvant CFA (Sigma Chemicals, St. Louis, MO, USA) supplemented with 4 mg/ml heat-killed *Mycobacterium tuberculosis* H37Ra (400 μg; Becton, Dickinson and Company, Sparks, MD, USA) as described previously [21-23]. One set of mice was injected with CFA without antigen to serve as a control. All the mice were housed with standard food and water ad libitum at a room temperature of 22 ± 2°C under a 12:12 h light-dark cycle. Clinical scores were monitored daily until the duration of the study in a blinded fashion by measuring paralysis according to the conventional grading system as described previously [21-23]. EAE mice treated with PBS or MaR1 were euthanized as indicated in the figures. SJL mice were anesthetized with CO_2_ and transcardially perfused with 1× chilled PBS, followed by perfusion with 4% paraformaldehyde for histopathological analysis and confocal imaging.

All the animal experiments in this study were performed as per the policies and guidelines of the IACUC committee at Henry Ford Health under the animal welfare assurance number D16-00090.

### MaR1 administration

The mice were randomly assigned to the treatment and control experimental groups (CFA, EAE, and EAE treated with MaR1; hereafter referred to as MaR1; N=8). The MaR1 solution was freshly prepared from the stock solution and administered immediately to prevent degradation. Daily intraperitoneal injections (i.p.) of 300 ng of MaR1 (7R,14S-dihydroxy-4Z, 8E, 10E,12Z, 16Z, or 19Z-DHA; Cayman Chemicals, Ann Arbor, MI, USA) per mouse in 200 µL of PBS or vehicle were given to each individual mouse beginning on day 6 after disease induction until the end of the study.

For the IL10-neutralizing experiments, SJL mice were immunized as described above (for EAE induction and functional evaluation), followed by i.p. injection of 200 µg of anti-IL10 (BioXCell # BE0049) twice a week and daily ip treatment with 300 ng of MaR1. Controls received isotype-matched rat IgG1 (BioXCell # BE0088) following the same protocol. The clinical symptoms were recorded as the clinical score, and the mice were subsequently euthanized at 21 days post immunization (dpi).

### Behavioral test

Locomotor activity was measured to assess the effect of MaR1 treatment on EAE. We employed an infrared (IR)-based technology called the novel infrared-based automated activity monitoring system (IRAMS) [24], developed by Columbus Instruments International (Columbus, OH). This instrument is supported by several electrodes embedded in an electric sensing board spaced 4 inches apart, with a beam diameter of 0.125 inches and a beam scan rate of 160 Hz. The sensors were properly screwed on a bracket to increase the flexibility and adjustability of the device. Acrylic cages were placed between the brackets. Locomotor activity in the form of diurnal horizontal (X-axis) and nocturnal vertical (Z-axis) activity was measured, and quantitative analysis was performed with MDI software (Columbus Instruments International, Columbus, Ohio) based on the system calculating the number of infrared interruptions emitted by mice at a specific time interval.

### Antigen Recall Response

For antigen recall response, spleen and lymph node (LN) cells (4×106/ml) were cultured in the presence or absence of the antigen PLP_139-151_ (20 μg/ml). The production of pro- and anti-inflammatory cytokines (IFNγ, GM-CSF, IL17a, IL10, and IL4) was examined as described previously [21-23, 25].

### Macrophage and CD4+ cell isolation

On day 18 postimmunization, F4/80 macrophages were isolated from the spleens of EAE mice treated with or without MaR1 using a Miltenyi Biotec Kit (#130-110-443; ∼92% purity). Similarly, CD4+ cells were isolated from the spleen using a CD4 isolation kit (BioLegend # 480033; ∼94% purity).

### Adoptive transfer

For adoptive transfer experiments, EAE was induced in SJL mice, and MaR1 treatment began when the mice in the randomly divided groups had recovered from disease. At the end of the study, lymph nodes were isolated from the MaR1-treated and untreated EAE groups and cultured with PLP_139-151_, anti-IFNy (10 µg/ml), IL12p70 and IL23 (10 ng/ml). After three days of incubation, 5 million CD4+ cells were injected intraperitoneally into SJL mice (n=5), followed by daily evaluation of the clinical score.

### Single Molecule Array (SIMoA) Assay

To assess the effect of MaR1 treatment on the CNS in the EAE groups, we measured the levels of neural markers (NFL and GFAP) in plasma samples from the MaR1-treated and untreated EAE groups with the commercially available SiMoA™ Neuro 2-Plex B Advantage Kit (product number: 103520) (Quanterix, MA) on an SR-X analyzer according to the manufacturer’s instructions. Additionally, the effect on inflammation was examined by cytokine profiling. We used a Mouse Cytokine 5-Plex Kit for IL6, TNFA, IL17, IL12, and IL10 (Cat # 107-178-1-AB; Product Number: 85-0441) (Quanterix, MA) on an SP-X analyzer according to the manufacturer’s instructions. We applied SIMOA, as this is an emerging ultrasensitive technology with the potential to detect analytes that are present at low levels and below detection limits for other conventional assays.

### RNA isolation and qRT−PCR

Total RNA was isolated from cells with QIAzol reagent using an RNeasy kit (Qiagen), and 1 µg of RNA was used for cDNA synthesis using an iScript cDNA synthesis kit (Bio-Rad) according to the manufacturer’s guidelines. Quantitative PCR analysis was performed using SYBR Green (Bio-Rad) on a Bio-Rad CFX96 real-time PCR detection system and software (CFX Maestro, Bio-Rad). Gene expression was normalized to that of the control housekeeping gene, ribosomal L27, using CFX Maestro software (Bio-Rad).

### Enzyme-linked immunosorbent assay (ELISA)

The protein levels of the cytokines IFN-γ, IL-17A, GMCSF, IL-4, and IL-10 were quantified using standard enzyme-linked immunosorbent assay (ELISA) kits provided by Biolegend and BD Biosciences.

### Immunohistology

Briefly, brain and spinal cord tissues were harvested from the euthanized groups, postfixed in 4% paraformaldehyde at 4°C for 48 h and cryoprotected with 30% sucrose until they sank. Tissues were subsequently embedded in OCT compound, and frozen sections (10-20 µm) were cut to assess the infiltration of immune cells and demyelination in the CNS. Histopathological evaluation via hematoxylin and eosin (H&E) and Luxol fast blue (LFB) staining methods was performed as described previously [23, 25]. Specifically, immunohistochemistry (IHC) was performed on CD4+ and F4/80+ cells from spinal cord samples to assess the effect of MaR1 treatment on the immune landscape inside the CNS. All images were captured by using a light microscope. For immunofluorescence staining, the sections were washed for 20 min with 0.1% PBS-Triton X-100, incubated with a blocking solution containing 1% BSA for 1 h at room temperature, and incubated with a mouse anti-MBP monoclonal antibody (1:500; Cell Signaling Technology) overnight at 4°C. Subsequently, the sections were washed and incubated with Alexa Fluor (AF) 568- or AF 488-coupled secondary antibodies (Life Technologies). Nuclei were counterstained with 4′,6-diamidino-2-phenylindole (DAPI) (H-1200; Vector Laboratories, Inc., CA, USA), and the cover slipped. Images were taken by a laser scanning confocal microscope (LSCM, Olympus). Images were quantified using ImageJ software (National Institutes of Health, Bethesda, MD).

### Flow cytometry

The immune cell profiles in the spleen/LNs and CNS tissues (brain and spinal cord) of EAE mice were determined at different time points for the corresponding experiments using standard staining methods and analysis protocols previously described in our published work [22, 23, 25]. All the data were processed using FlowJo® software and an analysis program (Treestar, Ashland, OR).

### Detection of reactive oxygen and nitrogen species (ROS and RNS)

The intracellular levels of ROS/RNS in different immune cells of the CNS were evaluated using the Cellular ROS/RNS Detection Assay Kit (Abcam, USA; catalog number: ab139473) according to the manufacturer’s instructions. Briefly, brain single-cell suspensions were prepared using a Percoll density gradient, and the cells were then treated with a mixture of two different fluorescent dye reagents in phenol-free culture media that allows the detection of ROS and RNS. After ROS/RNS dye treatment, the cells were washed, and Fc receptors were blocked with anti-mouse CD16/32 (BioLegend). The cells were then stained with fluorochrome-conjugated antibodies (anti-mouse-BV421-CD45) and anti-mouse-PE-O4 (Miltenyi Biotec)) for 40 min at 4°C to characterize oligodendrocytes (CD45^-^O4^+^). The cells were washed twice and resuspended in FACS staining buffer. The samples were analyzed on an Attune NxT cytometer (Thermo Fisher Scientific) using FlowJo software.

### Bioenergetics

To monitor the mitochondrial oxygen consumption rate (OCR) and extracellular acidification rate (ECAR) in intact cells, a Seahorse Bioanalyzer (Agilent) was used. The XFe mitochondrial stress test was performed on CD4+CD25-cells (purity ∼94%) or F4/80 macrophages (purity ∼92%) isolated from CFA or from EAE mice treated or not treated with MaR1 (300 ng/mouse daily) to measure the OCR. The cells were seeded at a concentration of 0.5 million per well (in the case of CD4+ cells) or 0.1 million per well (in the case of F4/80+) in a polylysine-coated XFe 96-well Seahorse culture microplate in 75 µl of OCR or ECAR DMEM and centrifuged at 2000 rpm for 1 min. Then, 100 µl of the respective media was added, and the mixture was incubated at 37°C in a CO_2_-free incubator for degassing. After incubation, the OCR and ECAR were measured according to the manufacturer’s protocol. To measure ATP levels, the cells were plated in 96-well plates (25,000 cells/well), and the ATP concentration was determined with a fluorometric ATP determination kit following the manufacturer’s protocol (Thermo Fisher). The data are presented as the unit/number of cells.

### Single Cell ENergetIc Metabolism by Profiling Translation inHibition (SCENITH)

We used the SCENITH assay, a FACS-based, quantitative analysis of mRNA translation in single cells [26-29]. It takes advantage of cells exposed to a short pulse of puromycin (PURO), which is incorporated into polypeptides that are being translated. The fixed cells were permeabilized and stained with an anti-PURO antibody. Translation is an energy-consuming process [30]. SCENITH is a proxy for the metabolic fitness of the cell, and its metabolic outcome correlates well with that of Seahorse [27, 28], requiring only one-tenth of the cell number. This approach provides information on the use of energetic pathways (glucose dependence, mitochondrial dependence, glycolytic capacity, and fatty acid and glutaminolysis capacity) [27, 29]. This method allowed us to analyze the metabolic activities of specific cell populations to examine the metabolic alterations in CD4+ T cells, macrophages, microglia, and O4+ oligodendrocytes using a previously described protocol and optimized assay conditions in our laboratory [29].

### Efferocytosis Assay

For the efferocytosis assay, FITC-labeled apoptotic splenic cells devoid of monocytes/macrophages were added to bone-derived macrophages (BMDMs) at a 1:5 ratio for 18 hours, followed by five washes with phosphate-buffered saline (PBS) to remove unbound apoptotic cells. The macrophages were then harvested with trypsin and centrifuged, and the pellet was mixed with FACS stain buffer. The macrophages were then stained with anti-mouse BV421-F4/80 for 30 minutes in the dark at 4°C. Flow cytometry was used to investigate the engulfment of apoptotic cells (FITC+) by BMDMs.

To demonstrate that BMDMs engulf apoptotic cells, macrophages were grown on a 4-chamber slide covered with polylysine, and FITC-labeled apoptotic splenic cells were then introduced for the same duration as indicated previously. After eliminating any unbound apoptotic cells, the BMDMs were washed five times with PBS. Then, the macrophages were fixed and permeabilized with 4% paraformaldehyde (PFA) and 0.1% Triton X-100. The BMDMs were then stained with phalloidin and DAPI and imaged using a confocal microscope.

### RNA sequencing

For transcriptome analysis, the spinal cords of three mice from each group, CFA-, EAE-, and MaR1-treated mice, were subjected to RNA-seq via the services of CD-Genomics, NY. The final library size was approximately 400 bp, and the insert size was approximately 250 bp. Illumina® 8-nt dual indices were used. Equimolar pooling of the libraries was performed based on the QC values, and the libraries were sequenced on an Illumina® NovaSeq 6000 (Illumina, California, USA) with a read length configuration of 150 PE for 40 M PE reads per sample (20 M in each direction). To assess the sequencing quality, a quality check (QC) was performed using FastQC during the data analysis. All the samples passed QC, which was performed using FastQC. Using the STAR aligner, paired-end reads of 150 bp were uniquely aligned to GRCm38-mm10, and a mapping rate of approximately 80% STAR was used to count the reads for each sample. The limma-Voom package in R was used to assess differential expression, with a padj cutoff of 0.05 and a logFC of 0.34. The threshold for filtering out genes with low expression was set to 1 CPM in at least 80% of the samples. The trimmed mean of M-values (TMM) method was used to normalize the counts. The significantly dysregulated genes were analyzed for Gene Ontology (GO) enrichment with an FDR cutoff of <0.05 using shinyGO.

### Peripheral blood cell isolation from MS patients and T-cell responses by flow cytometry

Peripheral blood mononuclear cells (PBMCs) were isolated after venous puncture from RR-MS patients (Table 1) and were separated by density gradient centrifugation over Ficoll-Hystopaque. PBMCs (1×10^6^ cells) were pretreated with or without MaR1 (10 nM) for 30 min and then stimulated with Dynabeads CD3/CD28 T-Cell Expander (one bead per cell; Invitrogen) for 8 h, as previously reported [31, 32]. To measure the intracellular cytokine levels, secretion was inhibited by the addition of 1 µg/ml brefeldin A (Sigma−Aldrich) 6 hours before the end of stimulation with either PMA/ionomycin or Dynabeads CD3/CD28 T-Cell Expander. At the end of the incubation, the cells were stained at the cell surface with Brilliant Violet 421-conjugated anti-CD3 (BioLegend), FITC-conjugated anti-CD4 (eBioscience), PerCP5.5-conjugated anti-CD8 (BioLegend), and made permeable with Cytofix/Cytoperm reagents (BD Biosciences) and then stained intracellularly with phycoerythrin-Cy7-conjugated anti-TNF-α (BD Biosciences), allophycocyanin (APC)-conjugated anti-IFN-γ (BD Biosciences) and phycoerythrin (PE)-conjugated anti-IL-17 (eBioscience) in 0.5% saponin at RT for 30 min. For Treg staining, the cells were stained at the cell surface with FITC-conjugated anti-CD4 (BD Biosciences), APC-conjugated anti-CD25 (BD Bioscience), fixed and permeabilized with the FoXP3 Transcription Buffer Kit (eBioscience) and intracellularly stained with PE-CF594-conjugated anti-Foxp3 (BD Biosciences) and PE-conjugated anti-IL-10 (BD Biosciences). All samples were acquired on a 13-color CytoFLEX flow cytometer (Beckman Coulter), and for each analysis, at least 300,000 events were acquired by gating on Pacific Orange-conjugated live/dead negative cells and analyzed by FlowJo Software.

**Table 1.** Demographic data of RR-MS patients.

|  | **RR-MS** |
| --- | --- |
| N° of subjects | n=8 |
| Mean age (years) | 34.10± 8.72 |
| Female/male | 5/3 |
| Disease duration* (years) | 6.4 |
| Relapse/Remission | 4/4 |
| Mean EDSS | 1.8 |
| DMT when sampling (yes/no) | 0/0 |
| Corticosteroids (yes/no)<br>(>1 month before sampling) | 0/0 |

## Data and statistical analysis

All values are presented as the mean ± SEM, and statistically significant differences were assessed by the Mann-Whitney test, Student’s t test, and one-way ANOVA. The statistical significance of the P values is described in the figure legends for each experiment. All the statistical analyses were performed with GraphPad Prism.

## Results

### MaR1 ameliorates clinical signs of EAE, improves pathophysiology, and promotes neuroprotection

To investigate the comprehensive protective mechanism of MaR1 in EAE, we induced EAE in SJL mice, a well-known relapsing-remitting animal model (RR-EAE) of MS. MaR1 treatment began after six days of immunization, while the other group received vehicle (0.1% ethanol). We found that mice treated with MaR1 exhibited significant protection against neurological impairments (**Fig. 1A-E**). EAE in SJL mice followed a classical relapsing-remitting pattern of disease (**Fig. 1B**). Compared with vehicle treatment, MaR1 treatment did not affect early-onset of disease; however, it significantly reduced disease severity at the peak stage compared to vehicle-treated EAE group (**Fig. 1B**). MaR1 treatment effectively prevented further relapses in the treated group (**Fig. 1B**). Compared with EAE vehicle-treated mice, MaR1-treated mice had significantly lower cumulative scores (p<0.01) and maximum scores (p<0.001) (**Fig. 1C**). Compared with vehicle treatment, MaR1 significantly reduced disease severity (p<0.001) and symptom incidence (p<0.0001) in EAE mice (**Fig. 1D-E**). Further, we measured the effect of MaR1 on the locomotor activity using the IRAMS monitoring system [24]. The average horizontal and vertical activity during the day was significantly lower in EAE mice than in animals treated with control (injected CFA lone) or MaR1. The MaR1 treatment increased the horizontal and vertical activity of treated group (**Fig. 1F**). Similar activities in mice were assessed after sunset, and we observed that horizontal and vertical activities in EAE mice were significantly lower than those in the CFA group, while MaR1-treated mice showed a significant improvement in activity compared to that of EAE mice (**Fig. 1G**). We also measured hourly changes in horizontal and vertical activity during the day and night. We discovered a significant decrease in average hourly activities in the EAE group compared to the CFA group, which was ameliorated by MaR1 treatment (**Fig. 1H**). Overall, we demonstrated that MaR1 treatment improved neurological function in EAE mice and prevented disease progression in the RR-EAE model.

**Figure 1:** Protective effect of MaR1 treatment on neurological deficits in the RR-EAE mouse model. **A**. Clinical scores of SJL mice in the CFA, EAE, and EAE groups treated with MaR1 for 70 days after disease induction. EAE mice developed disease symptoms beginning on day 9 after immunization with PLP, whereas MaR1-treated mice showed symptoms on day 10 (n=10). **B**. Cumulative score. **C**. Maximum score. **D**. Disease severity score. **E**. Incidence of clinical symptoms in EAE- and MaR1-treated mice. **p<0.01, ***p<0.001, ****p<0.0001 (as determined by the Mann−Whitney test) vs vehicle-induced EAE and MaR1 EAE. The data are shown as the mean ± SEM. For spontaneous motor activity, before the end of the experiment, locomotor activity of the mice was tested day time and overnight to evaluate disease severity. **F**. Average diurnal horizontal and vertical activities were measured in CFA-, EAE- and MaR1-treated mice. **G**. Average nocturnal horizontal and vertical activity were measured in CFA-, EAE- and MaR1-treated mice. **H**. Average hourly horizontal and vertical activity (diurnal and nocturnal) in CFA-, EAE- and MaR1-treated mice. The values are expressed as the means ± SDs (n=10). Statistical analyses were performed with one-way ANOVA and two-way ANOVA. *P<0.05, **P<0.01 versus the CFA group; #P<0.05, ##P<0.01 versus the MaR1-treated group.

### MaR1 reduces the antigen-specific immune response and immunomodulates infiltrating cells in the CNS

To investigate the effect of MaR1 on the myelin-specific immune response, we isolated splenic and lymph node (LN) cells from both the EAE and MaR1-treated groups and stimulated them with PLP_139-151_. After 72 hours of stimulation, the cell supernatant was tested for pro- and anti-inflammatory cytokines. **Figure 2A** shows that MaR1 treatment decreased antigen-induced IL17a production in spleen/LN cells while increasing the expression of IFN*γ* and anti-inflammatory cytokines, such as IL4 and IL10. This observation was supported by the results of quantitative PCR (qPCR) analysis of pro- and anti-inflammatory cytokine levels in spleen/LN cells (**Supp. Fig. 1A-B)**. These findings indicate that MaR1 treatment alters the antigen-specific immune response.

**Figure 2:** MaR1 modulates antigen-specific responses and abrogates the infiltration of IL17- and IFNg-producing CD4+ T cells. **A**. The antigen recall response of spleen/LN cells was examined on day 17 after immunization for 72 h in the presence of PLP_139-151_ (n=4). **B-D**. CNS tissues (brain and spinal cord both) were processed, and the total number of leukocytes (CD45+) and CD4+ T cells and the frequencies of Th1 and Th17 cells and myeloid cells in the CNS of treated and untreated RR-EAE (n=4) were examined. **E**. Infiltrating myeloid cells, including monocytic DCs, F4/80+ macrophages and monocytes, were profiled in both groups (n=4). **F-G**. At the peak of the disease, proinflammatory (class II and CD38) and anti-inflammatory (EGR2 and CD206) effects on splenic macrophages (CD11b^+^F4/80^+^) were examined by flow cytometry, and the data are presented as a bar graph of the MFI (n=4). **H**. The ratio of CD206+/CD38+ macrophages was plotted to determine the macrophage phenotype (n=4). **I**. The level of neurofilament-light chain (NFL) in the plasma of EAE model mice treated with or without MaR1 was examined using SIMOA (n=5). **J-K**. Spinal cord sections showing inflammatory infiltrates (H&E) and demyelination (LFB). Representative images showing histopathological changes in spinal cord tissues from EAE mice treated with MaR1 or vehicle. The data are shown as the mean ± SEM. *, P<0.001; ***, P<0.01; **, P<0.05*.

Since T cells and myeloid cells contribute to autoimmune CNS inflammation and demyelination [33], we used flow cytometry to analyze the infiltrating cells in the CNS. We found that MaR1 treatment greatly reduced the infiltration of leukocytes (CD45+) and CD4+ cells (CD45+CD4+) into the CNS (**Fig 2C-D**). Because CD4+ T cells play an important role in EAE disease pathogenesis, we measured the total number of infiltrating CD4+ Th subsets, including Th1 and Th17 subsets, in the CNS tissues (brain and spinal cord) of the vehicle- and MaR1-treated EAE groups. MaR1 treatment significantly decreased the number of IFNγ-expressing (Th1) and IL17-expressing (Th17) CD4+ T cells in the CNS tissues of treated EAE mice (**Fig. 2E**), indicating that pathogenic CD4+ T cells are impacted by this proresolving mediator. We also examined the effect of MaR1 treatment on myeloid cell infiltration and found that MaR1 treatment significantly reduced the number of monocytic dendritic cells (CD45+CD11b+CD11c+MHC II+) and macrophages (CD45+CD11b+F4/80+) but had no effect on the number of infiltrated monocytes (CD45+CD11b+Ly6G-Ly6C+) (**Fig. 2E**). Monocytes/macrophages are among the major effectors of demyelination in both MS and EAE and are highly plastic in nature [34, 35]. Depending on the environment, monocytes differentiate into either proinflammatory or anti-inflammatory macrophages, and their ratio determines the outcome of the disease [23]. Since anti-inflammatory macrophages ameliorate the clinical symptoms of EAE [23, 36], we examined the splenic nature of macrophages in the MaR1-treated and untreated EAE groups by investigating the expression of the anti-inflammatory markers CD206 and EGR2 [37, 38] and the proinflammatory markers MHC-II and CD38 [38]. Notably, MaR1 treatment not only reduced MHC-II and CD38 expression and concomitantly induced EGR2 and CD206 expression (**Figs. 2F-G**) but also increased the EGR2/CD38+ and CD206/CD38+ macrophage ratios in the treated group compared to those in the EAE group (**Fig. 2H, Supp. Fig. 1C**), suggesting that MaR1 promotes a shift in macrophage polarization.

Blood neurofilament light chain (NFL) is a neuronal damage marker that has been linked to relapses, deterioration of the EDSS score, lesions on MRI images, and atrophy of both the brain and spinal cord in MS patients [39-41]. Using very sensitive single-molecule array technology (SiMoA), we found that MaR1 treatment significantly reduced NFL levels compared to those in the EAE group (**Fig. 2I**). The clinical score of MaR1-treated EAE mice was reduced, which was evident from the decrease in the number of infiltrating leukocytes and the preservation of myelin content in the spinal cord sections, as observed through hematoxylin and eosin and Luxol fast blue staining, respectively (**Figs. 2J-L**). Based on the histopathological analyses, MaR1 treatment significantly reduced the number of immune cells in mice with EAE compared to that in mice treated with vehicle (**Fig. 2J**). Furthermore, the occurrence of demyelination was considerably lower in the group of mice treated with MaR1 than in the group of vehicle-treated EAE mice (**Fig. 2L**). Overall, MaR1 effectively inhibited EAE disease progression by reducing the infiltration of immune cells into the central nervous system (CNS), blocking Th1 and Th17 immune responses, polarizing macrophages toward an anti-inflammatory phenotype and providing protection against neuroaxonal injury and myelin loss.

### MaR1 shows therapeutic potential in the RR-EAE model by impeding the encephalitogenic property of CD4 effector cells

To investigate the therapeutic effect of MaR1 on EAE, we treated RR-EAE animals in remission on day 20 after immunization, when all of the mice had recovered from paralysis, and the clinical score was investigated during subsequent relapse (30-38 dpi). We observed that MaR1 treatment significantly reduced relapse in RR-EAE mice, with animals showing a clinical score close to 0 (**Fig. 3A**). To better study how MaR1 affects CD4 T-cell effector function, we isolated cells from lymph nodes on day 38 and cultured them under Th17-inducing conditions with PLP_139-151_ (20 µg/ml) (**Fig. 3B**). After 3 days, CD4+ T cells were isolated and adoptively transferred (AdT) into naïve SJL mice, and the clinical score was monitored daily. CD4+ T cells obtained from EAE mice were able to induce EAE, while those from the MaR1-treated group were incapable of inducing EAE (p<0.01). These findings indicate that MaR1 has therapeutic potential and that treatment affects the encephalitogenic properties of effector CD4+ T cells.

**Figure 3:** MaR1 treatment impedes the encephalitogenic property of CD4 effector cells. **(A)** EAE was induced in SJL mice, and MaR1 treatment began when mice in the randomly divided groups had recovered from disease on day 20 postimmunization. **(B)** At the end of the study (∼38 days), isolated lymph node cells from both groups were cultured with PLP_139-151_, anti-IFNy (10 µg/ml), IL12p70 and IL23 (10 ng/ml). After three days, the enriched CD4+ cells were injected into SJL mice (n=5), after which the clinical score was monitored. <u>Note</u>: The Adt-EAE-MaR1 group was not treated with MaR1. P<0.01**.

### MaR1 mediates its protection by producing IL10 in EAE

Because MaR1 promotes the expression and production of IL10, which can influence Th17 pathogenicity in autoimmune disorders [42], we next investigated the immune cells that produce IL10 in response to MaR1 treatment. To do this, spleen cells from the vehicle- and MaR1-treated EAE groups were stimulated with PMA/ionomycin, followed by intracellular IL10 staining and surface staining for various immune cells. MaR1 treatment dramatically increased the percentage of total IL10-producing cells (p<0.01) (**Fig. 4A**). Treatment also considerably enhanced IL10 expression in different IL-10-expressing cell types, including CD4+ cells, CD8+ cells, B cells, and dendritic cells (DCs), although the greatest contribution was derived from CD4+ T cells and macrophages (∼4.5-fold) (**Fig. 4B and Supp Fig. 2**). Furthermore, compared with vehicle-treated EAE mice, MaR1-treated EAE mice had significantly greater plasma levels of IL-10 (p<0.01) (**Fig. 4C**).

**Figure 4:** Effect of IL10 neutralization on the protective effect of MaR1 on EAE. **(A)** Spleen cells from RR-EAE mice treated with or without MaR1 (day 18) were stimulated with PMA/ionomycin in the presence of GolgiPlug for 4 h and subjected to surface staining for CD4 (CD3+CD4+), CD8 (CD3+CD4+), B cells (CD3-CD19+), mDCs (CD11b+CD11c+Class II+) and macrophages (F4/80+) and intracellular staining for IL10. **(B)** CD45+IL10+ cells were gated from live cells; based on surface markers, all cell types expressing IL10 were profiled (n=4). **(C)** Plasma levels of IL-10 were measured using SIMOA on day 70 in the treated and untreated EAE groups treated with MaR1 (n=4). **D**. Disease severity plot showing abrogation of the protective effect of MaR1 by an IL-10 neutralizing antibody compared to that of an IgG neutralizing antibody (n=8). **E i-ii**. The number of CNS-infiltrating Th17 cells in all groups was determined via flow cytometry; the data are presented as a bar graph (n=5). **F**. Adoptive transfer of antigen-specific CD4+ T cells derived from the IL10 neutralization experimental batch and monitoring of clinical scores (n = 5). *p<0.05 vs. vehicle EAE. The data are shown as the mean ± SEM. p<0.01**, p<0.001*** compared to the EAE group.

To test whether MaR1-induced IL-10 was responsible for the observed protective effects exerted by this proresolving mediator, we neutralized IL-10 with an anti-IL-10 neutralizing antibody and found that, compared to IgG as a control, the anti-IL-10 antibody counteracted the protective effects of MaR1 in terms of the clinical score (**Fig. 4D**) and increased the infiltration of IL17-producing CD4+ T cells into the CNS (**Fig. 4E i-ii**). Furthermore, to investigate whether IL10 neutralization could affect MaR1-induced loss of encephalitogenic characteristics in CD4+ cells, LN cells obtained at 21 dpi were grown under Th17 conditions as reported in **figure 4B**, and on day 3, CD4+ T cells were adoptively transferred into naïve SJL mice, and the clinical score was assessed daily. As shown in **Figure 4FC**, CD4+ T cells from MaR1+aIL10-treated animals restored their encephalitogenic properties to levels comparable to those of the EAE-Adt group. Taken together, these findings confirm that MaR1 mediates its protective effect via IL10.

### MaR1 promotes metabolic reprogramming in effector CD4+ T cells and macrophages

Since metabolic reprogramming is crucial for T-cell activation, differentiation, and function [43, 44], we next sought to investigate whether MaR1 has an impact on CD4 T-cell metabolism *in vivo*. CD4+ T cells (CD4+CD25-) isolated from MaR1-treated mice exhibited significantly greater spare respiratory capacity than CD4+ T cells from EAE mice, which exhibited decreased mitochondrial respiration (**Fig. 5A**). MaR1 improved the glycolytic rate in CD4+ T cells (**Fig. 5B**), resulting in a ‘higher metabolic state’, whereas vehicle-treated EAE CD4+ T cells had a ‘lower metabolic state’ (**Fig. 5C**); however, no change in total cellular ATP levels was observed (**Fig. 5D**). These results suggest that MaR1 treatment affects the metabolic and bioenergetic states of CD4+ T cells *in vivo*.

**Figure 5:** MaR1 induces metabolic reprogramming in CD4+ T cells and macrophages in EAE mice. **A**. An XF mitochondrial stress test was performed on CD4+CD25-cells (purity ∼95%) isolated from CFA-, EAE- and MaR1-treated EAE mice on day 18 postimmunization. The maximal respiration is presented as a bar graph (n=6). **B**. Compensatory glycolysis was examined using an XF Seahorse bioanalyzer and is presented as a bar graph (n=6). **C**. The bioenergetic profile depicts the metabolic state of CD4+ cells isolated from the various groups described in A. **D**. ATP levels detected in CD4+ cells from the various groups in A using an ATP assay kit (n = 4). NS, not significant compared to the control and EAE groups; Student’s t test was used. **E**. At the peak of the disease, brain infiltrating cells (BILs) were isolated from all groups using a Percoll gradient and processed for SCENITH. MFI of puromycin across samples treated with different inhibitors of CD4+ T cells (CD45^+^CD4^+^), including deoxyglucose (DG), oligomycin (OM), or deoxyglucose + oligomycin (DGO) (n=3). The metabolic perturbations in the infiltrated CD4+ cells in the various groups are shown as a bar graph. **F-G**. An XF mitochondrial stress test was performed on splenic F4/80+ macrophages (purity ∼95%) isolated from CFA, EAE and EAE mice treated with MaR1. The maximal respiration, basal glycolysis, bioenergetic profile and total ATP levels were detected in macrophages from the various groups as described above in detail. **H**. The metabolic perturbation of the infiltrated macrophages (CD45^+^CD11b^+^F4/80^+^) in the various groups was detected using SCENITH as described above. The data are shown as the mean ± SEM (n = 3). NS, not significant; ***p < 0.001 compared to the control or EAE group using Student’s t test.

To further investigate the effect of MaR1 on the metabolic fitness of infiltrated CD4+ T cells in the CNS, we used Single Cell ENergetIc Metabolism by Profiling Translation inHibition (SCENITH), which allows us to assess the metabolic state of numerous cell types by flow cytometry according to the gating strategy shown in Supplementary Fig. 3. To do so, infiltrating mononuclear cells were isolated at 18 dpi, and single cells were treated with oligomycin (OM), 2DG, or both (OM+2DG), followed by a brief pulse of puromycin (PURO), allowing us to quantify metabolic fitness in terms of mitochondrial dependency, fatty acid and amino acid oxidation (FAO & AAO) capacity, and glycolytic and glucose dependence [27, 29]. We found that infiltrated CD4 T cells from the MaR1-treated groups had significantly greater mitochondrial dependency, FAO, and AAO capability than those from the EAE group (**Fig 5E**). Furthermore, MaR1 reduced the glucose reliance of infiltrating CD4+ T cells without impacting glycolysis compared to that of infiltrating CD4+ T cells from EAE.

Given that MaR1 treatment induced macrophage polarization toward an anti-inflammatory phenotype (**Fig. 2F-H**) and that metabolic reprogramming plays a major role in macrophage phenotypic alterations [23, 45], we investigated the metabolic status of F4/80+ macrophages isolated from EAE mice treated with or without MaR1 at 18 dpi and found that MaR1 treatment significantly decreased mitochondrial respiration and glycolysis compared to those of macrophages isolated from the CFA group, and these effects were reversed by MaR1 treatment (**Figs. 5F-G**). Interestingly, macrophages from the vehicle-treated group exhibited a greater metabolic state, whereas EAE macrophages exhibited a lower metabolic state, which was reversed by MaR1 treatment (**Fig. 5H**). Although there was no difference in total cellular ATP levels between the EAE and CFA groups, MaR1 significantly increased ATP levels between the CFA and EAE groups (**Fig. 5I**), implying that an increased energetic state in macrophages in the MaR1 group may have contributed to energy production. Similar to CD4+ T cells, we investigated the metabolic fitness of F4/80 macrophages infiltrated by SCENITH according to the gating strategy shown in Supplementary Fig. 3 and found that macrophages in the MaR1-treated group had significantly greater mitochondrial dependency, FAO and AAO capability, and lower glucose and glycolytic dependence than those in the vehicle-treated EAE group (**Fig. 5J**). Overall, these results indicate that MaR1 treatment shapes the metabolic and bioenergetic states of both peripheral and infiltrating CD4+ T lymphocytes and macrophages.

### MaR1 enhances macrophage efferocytosis

Since removal of dead cells by macrophages, known as “efferocytosis,” is a key proresolving functional test [46-48] and many neurodegenerative disorders are caused by efferocytosis dysfunction[49], flow cytometry and confocal imaging were used to analyze the efferocytosis ability of these cells and revealed that MaR1-treated macrophages exhibited greater efferocytosis than did vehicle-treated macrophages (**Fig. 6A**). Under efferocytosis conditions, MaR1 treatment led to increased expression of the anti-inflammatory cytokines IL10 and TGFβ (**Fig. 6B**). Furthermore, we observed a similar impact of MaR1 *in vivo* by analyzing brain sections, and we observed that spinal cord CD68+ macrophages from the MaR1-treated EAE group displayed a greater phagocytic index for myelin debris than those from the vehicle-treated EAE group (**Fig. 6C**). These findings show that MaR1 promoted macrophage efferocytosis both *in vitro* and *in vivo*.

**Figure 6:** MaR1 induces efferocytosis in macrophages. **Ai**. Bone marrow-derived macrophages were treated with MaR1 (100 nM) or vehicle (0.1% EtOH) for 1 h, after which CFSE-labeled 70-75% apoptotic splenic cells devoid of monocytes/macrophages were added at a ratio of 1:5. After 18 h, the cells were washed and stained for F4/80, after which the number of F4/80 cells engulfing CFSE-labeled cells was quantified (n=6). The data are presented as a bar graph and a representative flow plot. **Aii**. Another set of cells was plated on coverslips, and an *in vitro* efferocytosis assay was performed as described above. After 18 h, the cells were washed and fixed, and Z-stack images were taken with an Olympus FV1000 confocal microscope with a 40× objective. Four consecutive images with an interspace of 1 µm were captured. The mean intensity of apoptotic cells was quantified with ImageJ analysis software (version 1.49; NIH, USA) (n=6). **B**. Under efferocytosis conditions, the expression of IL10 and TGFβ was examined by qPCR, and the data were normalized to those of the housekeeping gene L27 (n=3). **C**. For efferocytosis in the spinal cords of the EAE and MaR1-treated groups, macrophages (CD68+) were stained with a mouse anti-CD68 antibody and polyclonal degraded myelin (Millipore). The images were captured by LSCM at a 1-ary unit aperture at 60x resolution. P<0.001*** compared to Wt.

### MaR1 induces metabolic reprogramming in microglia by switching them to an anti-inflammatory state

Microglia, or brain-resident macrophages, play dual roles in neuroinflammation [50]. Depending on the environment, microglia can be either protective or detrimental to neurodegeneration, polarizing them into pro-or anti-inflammatory states. In our study, MaR1 treatment decreased the expression of the proinflammatory markers MHC-II and CD38 while increasing the expression of the anti-inflammatory markers EGR2 and CD206 (**Fig. 7A**). The ratios of pro-vs. anti-inflammatory markers (EGR2/CD38 and CD206/Class II) (**Fig. 7B**) demonstrated that MaR1 promotes a switch toward an anti-inflammatory phenotype. Similar to macrophages, MaR1 also significantly induced efferocytosis in primary microglia (**Fig. 7C, Supp. Fig. 4**), as microglia are also professional CNS phagocytes that engulf myelin debris and initiate synaptogenesis and neurogenesis [51]. We recently used an EAE model to study the metabolic control of microglia during neuroinflammation *in vitro* and *in vivo* [29]. Thus, we investigated whether MaR1 treatment restored metabolic perturbations in microglia in the EAE group. Similar to our previous observation [29], utilizing SCENITH and flow cytometry (Supplementary Figure 3), we observed that microglia in the EAE group had lower mitochondrial function and greater glycolysis than those in the CFA control group, which was reversed by MaR1 treatment (**Figure 7D**). These findings demonstrate that MaR1 induces an anti-inflammatory phenotype in microglia, which may contribute to the proresolving effects of MaR1 on EAE via metabolic reprogramming.

**Figure 7:** MaR1 promotes an anti-inflammatory phenotype in microglia. **A**. At the peak of the disease at ∼18 days, BILs from all groups (EAE- and MaR1-treated) were processed for pro- and anti-inflammatory marker detection by flow cytometry (n = 3). **B**. Bar graph of the EGR2/CD38 and CD206/Class II ratios. **C**. Primary microglia were treated with MaR1 (100 nM) or vehicle (0.1% EtOH) for 1 h, after which CFSE-labeled 70-75% apoptotic splenic cells devoid of monocytes/macrophages were added at a ratio of 1:5. After 18 h, the cells were washed and stained for CD11b, after which the number of CD11b+ microglia engulfing CFSE-labeled cells was quantified (n=3). The data are presented as a bar graph. **D**. Changes in the metabolism of microglia in the various groups were evaluated using SCENITH, and the data are presented as a bar graph. The data are shown as the mean ± SEM (n = 3). NS, not significant; ** < 0.05; **p < 0.01; **** or ***p < 0.001 compared to CFA or EAE.

### MaR1 promotes oligodendrocyte metabolic fitness during EAE

Since MaR1 was able to decrease myelin loss, as shown by LFB staining (Fig. 4C), we next investigated whether this effect was due to the active role of oligodendrocytes. Representative immunofluorescence images confirmed greater myelin basic protein (MBP) expression in the white matter (carpus callosum) of the brain of the MaR1-treated group than in that of the untreated EAE group (**Fig. 8A**). Furthermore, MaR1 treatment reduced the levels of reactive oxygen species (ROS) and reactive nitrogen species (RNS) in O4+ oligodendrocyte populations (**Fig. 8B**). The metabolic parameters of these cells were investigated using SCENITH, flow cytometry and PURO MFI (**Fig. 8C, Supplementary Fig. 3**), and we found that O4+ oligodendrocytes from EAE mice were glycolysis dependent and exhibited a significant reduction in mitochondrial respiration and fatty acid oxidation (FAO), which were restored by MaR1 treatment (**Fig. 8D**). These findings suggest that MaR1 may protect against demyelination and restore dysfunctional oligodendrocyte metabolism.

**Figure 8:** MaR1 protects myelin possibly by regulating the metabolic fitness of oligodendrocytes during EAE. **A**. Serial coronal brain sections from CFA-, EAE- and MaR1-treated EAE mice on day 18 post immunization were stained with fluoromyelin, and imaging was performed using laser scanning confocal microscopy. The intensity of myelin in the CC above the lateral ventricle was measured with ImageJ (labeled LV) (n=3). **B**. At the peak of the disease, single suspensions of CNS tissues were processed using a Percoll gradient from all groups (CFA, EAE and MaR1 treated), and isolated cells were processed for the detection of intracellular reactive oxygen and nitrogen species (ROS and RNS) levels in O4+ oligodendrocytes (CD45^-^O4^+^), which were evaluated using the Cellular ROS/RNS Detection Assay Kit by flow cytometry (n = 5). **C**. Cellular metabolism of O4+ oligodendrocytes (CD45^-^ O4^+^) was examined using SCENITH (n=3). The metabolic changes, including changes in mitochondria, glycolysis and fatty acid oxidation (FAO), in O4^+^ oligodendrocytes in the various groups are shown as a bar graph. *p<0.05, **p<0.01, ****or ***p< 0.001 compared to CFA or EAE.

### MaR1 reverses EAE-induced alterations in the spinal cord transcriptome

Since MaR1 exerts several protective effects at different levels and involving different cellular types, we investigated whether transcriptome changes were associated with MaR1 treatment by performing RNA-seq on the spinal cords of the CFA, EAE, and MaR1 groups on day 70 dpi, namely, after the relapsing-remitting course of the disease. Interestingly, we found a clear distinction between the groups (**Fig. 9A**), with numerous differentially expressed genes. Across the groups, 631 genes were significantly upregulated in the EAE group but downregulated in the MaR1 treatment group (**Fig. 9B-C**). By investigating the functional relationships of the dysregulated genes using Gene Ontology (GO) analysis, the downregulated genes were found to be associated with the inflammatory response, leukocyte activation and other immune/defense-related pathways (**Fig. 9D**). Furthermore, MaR1 dramatically increased the expression of a limited subset of genes involved in axonogenesis and related functions, the majority of which were downregulated in EAE (**Fig. 9E**). Notably, MaR1 did not have any upregulated or downregulated genes in the same direction as EAE, suggesting that MaR1 helps prevent EAE-induced inflammation and may even concomitantly drive axonal regeneration.

**Figure 9:** MaR1 rescues EAE-induced spinal cord transcriptomic alterations. **(A)** MDS plot showing the segregation of each group: CFA, EAE and MaR1. **(B)** Heatmap representing significant gene expression changes across the groups. Padj >0.05. (**C)** Venn diagram representing the shared number of up- and downregulated genes in the EAE+MaR1 group. (**D)** Gene Ontology (GO) enrichment of the common genes upregulated with EAE and downregulated by MaR1 treatment. FDR < 0.05. **(E)** Gene Ontology (GO) enrichment of the common genes downregulated with EAE and upregulated by MaR1 treatment. FDR < 0.05.

### MaR1 reduces T-cell autoreactive responses and promotes IL10-producing Tregs in RRMS patients

To translate these findings to a human setting, we next investigated whether MaR1 could also impact human T-cell responses. For this purpose, we investigated human T cells from patients with relapsing-remitting MS (**Table 1**). In particular, the immunomodulatory role of MaR1 was investigated upon polyclonal activation of the T-cell receptor with anti-CD3 and anti-CD28 antibodies, and the effect of MaR1 on specific T-cell subsets was investigated in CD4+ Th1 and Th17 cells and cytotoxic CD8+ T cells. As expected, and as previously shown[32], anti-CD3 and anti-CD28 stimulation resulted in high percentages of TNF-α-, IFN-γ-, and IL-17-producing T cells in all T-cell subsets (**Figs. 10B-C**). MaR1 treatment significantly impacted Th1 and Th17 responses by significantly halting IFN-γ and IL-17 production (**Fig. 10B**) and cytotoxic CD8+ T cells by reducing TNF-α and IFN-γ production (**Fig. 10C**). Moreover, MaR1 concomitantly potentiated Treg responses by significantly enhancing the expression of the signature transcription factor FoxP3 and their production of IL-10 (**Figs. 10D-E**).

**Figure 10:** MaR1 reduces cytokine responses in activated human T-cell subsets in patients with relapsing-remitting (RR)-MS. **A**. Peripheral blood mononuclear cells (1 × 10^6^ cells per well) were left untreated or treated with vehicle or MaR1 (10 nM) for 30 min (n = 8). The cells were then stimulated with anti-CD3/CD28 for 8 h, stained at the cell surface and intracellularly, and analyzed by flow cytometry. **B-C**. Cytofluorimetric plots and percentages of intracellular IFN-γ and IL-17 produced by CD4^+^ T cells and of TNF-α and IFN-γ produced by CD8^+^ T cells. **D-E**. Cytofluorometric plots and percentages of intracellular FoxP3 and IL-10 in Tregs. NS not significant, *p<0.05, **p<0.01, ****or ***p< 0.001.

## Discussion

Despite the availability of numerous immunomodulatory drugs, there are no entirely satisfactory treatments for MS, as most drugs have substantial side effects with less-than-optimal tolerance and do not provide complete remission or stabilization of disease. Given that unresolved inflammation is a hallmark of MS pathogenesis, it is imperative to study the endogenous mechanisms that can curtail excessive inflammation and promote its timely resolution. In MS, disease progression might partly be a consequence of failed inflammation resolution associated with metabolic dysfunction mediated by resolution mediators [12]. However, there is a gap in knowledge about the role of endogenous resolution mediators in MS. Identifying a therapeutic option that can modulate adaptive and innate immune responses without generally suppressing the immune system has been a key barrier to improving the treatment of patients with MS. Using metabolic profiling, we have shown in our previous work that the precursor of MaR1 (14-HDHA), a proresolving lipid metabolite of omega-3 polyunsaturated fatty acids, is significantly lower in the plasma of patients with MS [13].

A recent study showed that MaR1 has a protective effect on a chronic model of EAE [52]. However, its impact on RR-EAE and the underlying mechanisms have not yet been reported. In this study, we investigated the impact of MaR1 on inflammation and disease severity in an RR-EAE model and investigated how MaR1 treatment polarized adaptive and innate immune cells to dampen the antigen-specific immune response in this RR-EAE model. Our findings indicate that administering MaR1 at a dosage of 300 ng/mouse on day 6 after immunization or during the remission of clinical symptoms significantly reduced the severity of symptoms in the EAE model group. The studies presented here provide evidence for the potential of maresins for use in clinical settings. This is particularly significant because maresins do not suppress the immune system [53], unlike steroids and many other treatments for multiple sclerosis (MS). These findings suggest that MaR1 could be a new and powerful therapeutic alternative for MS that does not have immunosuppressive effects. Nevertheless, the administration of MaR1 reduced the production of IL17 and stimulated the production of IL4 and IL10. These results emphasize the immunomodulatory function of MaR1 in EAE by reducing proinflammatory conditions in the central nervous system and adjusting T-cell responses to promote an anti-inflammatory state. However, the beneficial effects of MaR1 on reducing neuroinflammation in EAE are substantial, as MaR1 significantly protects against neurological impairment and myelin loss, even when treatment is initiated on day 6 after immunization. In summary, our findings indicate that MaR1 expedited the resolution of inflammation in EAE and effectively halted the course of the disease. In addition, MaR1 has been implicated in various disease models, such as spinal cord injury [15], spinal muscular atrophy [54] and cerebral ischemia [17]. The proresolving role of MaR1 in macrophages is facilitated by its interaction with its receptor, G-protein coupled receptor 6 (LGR6) [55]. However, its impact on CD4+ cells has not yet been studied. LGR6 expression has been observed in regulatory T cells and has been found to facilitate the protective impact of MaR1 in a model of respiratory viral infection [56]. Nevertheless, the process by which the MaR1-LGR6 cascade influences the progression of EAE disease has not yet been examined.

Several studies have demonstrated the involvement of DHA in protecting the brain, but the specific processes involved are not yet fully understood. These mechanisms may include the maintenance of the myelin sheath, which helps to preserve the integrity of axons and hence facilitates the process of remyelination in brain diseases [57, 58]. DHA-derived metabolites possess neuritogenic, synaptogenic, and neurogenic characteristics. Studies have demonstrated that treatment with DHA itself and certain metabolites produced from it can improve or resolve inflammation, thereby serving as a protective nutritional shield for the brain [57, 59-64]. In the past, our group reported the very first study of the therapeutic effects of a specific SPM called resolvin D1 (RvD1) on the progression of disease in a preclinical animal model of MS using two different models of EAE [22]. We reported that RvD1 suppressed EAE disease progression by suppressing autoreactive T cells and inducing an anti-inflammatory phenotype in macrophages [22]. In addition, a recent publication provided further evidence of the beneficial effects of DHA metabolites (RvD1 and protectin D1) in preventing inflammation-induced dysfunction of the blood-brain barrier (BBB) [14]. Together, these studies offer vital information regarding the therapeutic capacity of metabolites produced from DHA for the treatment of MS.

Our research revealed that MaR1 functions as an immunomodulator, as its treatment caused changes in the immune system of EAE mice within the central nervous system. Unlike the results reported by Sánchez Fernández *et al*. [52], MaR1 significantly increased the RNA and protein levels of anti-inflammatory cytokines, specifically IL4 and IL10. This led to the promotion of an anti-inflammatory state in CD4 T cells and a decrease in the number of pathogenic CD4+ cells that produce IL17. IL10 plays a crucial role in the pathogenesis of multiple sclerosis (MS) and other neurological illnesses [65]. Consequently, any molecule, such as MaR1, that may increase the production of IL10 has significant promise for therapeutic advancement.

Efferocytosis is crucial for the recovery of homeostasis of the brain parenchyma, not only because it removes apoptotic cells before they progress to secondary necrosis, releasing toxic intracellular compounds and autoantigens but also because it modulates the phagocyte inflammatory response [66, 67]. Efferocytes are mainly monocytes/macrophages and brain-resident macrophages (microglia) [68]. Failed efferocytosis is emerging as a key mechanism driving the progression of chronic inflammatory diseases, including metabolic diseases and neurodegenerative diseases [49, 69]. It is impaired in several pathophysiological processes, giving rise to chronic inflammatory and neurodegenerative diseases such as MS [70, 71]. CNS diseases share some cardinal neuropathological events, such as inflammation and excitotoxicity, thus raising the possibility of phagocytosis impairment during the disease course [72, 73]. There is a lack of understanding of how efferocytosis plays a protective role during MS. Efferocytosis relies on metabolic signaling [74, 75] to stimulate anti-inflammatory responses, and pharmacological manipulation of metabolic pathways can be critical for promoting repair and healing in the body [76]. We observed that MaR1 restored metabolic reprogramming in macrophages/microglia, which could be one of the potential mechanisms of MaR1-mediated reversal of impaired efferocytosis in EAE. Moreover, its treatment induced the production of IL10 and TGFbeta, which are anti-inflammatory cytokines that are protective against EAE, under efferocytosis conditions. Additional in-depth studies are required to elucidate the mechanism by which MaR1 controls efferocytosis in EAE models. This could reveal new concepts and unexplored possibilities for the development of innovative treatments that enhance the healing and rejuvenation of the central nervous system not only for MS but also for other neurological disorders.

MaR1 can also protect microglia by supporting an anti-inflammatory state and initiating metabolic reprogramming. This indicates that microglia might play a role in restoring the CNS microenvironment toward the resolution phase in EAE. A number of studies have reported the protective role of microglia against EAE pathology [77-79]. Microglia also play a critical role in remyelination processes [80]. An increase in the anti-inflammatory phenotype of microglia and myelin protection in MaR1-treated EAE mice suggested the possibility of interplay between microglia and oligodendrocytes. Importantly, MaR1 maintains myelin status by inducing metabolic changes and reducing oxidative stress in oligodendrocytes, myelin-producing cells in the CNS, suggesting that the neuroprotective effects of MaR1 are partly induced by metabolic reprogramming of disease-associated cell types. Abnormal metabolic pathways in oligodendrocytes have been linked to a variety of neurodegenerative disorders [81, 82]. We and others have demonstrated that metabolism is critical for the maturation of OPCs into preoligodendrocytes [83] and for the shift from preoligodendrocytes to mature oligodendrocytes [84]. Given the critical role of oligodendrocytes in remyelination, limited information is available concerning MaR1-induced metabolic changes during oligodendrocyte development and how these changes contribute to the maintenance of myelination status. However, further studies are needed to establish whether MaR1 treatment provides neuroprotection or has a direct effect on oligodendrocytes progenitor cells to induce remyelination.

## Conclusion

Our data demonstrate that MaR1 exerts a broad impact on the functional capabilities of immune cells associated with EAE and MS, resulting in decreased severity of the disease, prevention of its progression, and improved neurological outcomes in the RR-EAE model of multiple sclerosis (MS). However, further studies are needed to determine the role of MaR1 in regulating the metabolism of these cells and how it impacts their effector functions and disease outcomes.

## Availability of data and materials

All data generated or analyzed during this study are included in this article and its supplementary information files.

## Declarations

### Ethics Approval

The animal studies performed in this manuscript were approved by the IACUC committee of Henry Ford Health. All human subjects provided informed written consent for the study, which was approved by the ethics committee of IRCCS Neuromed (protocol 26.03.2018).

## Consent for publication

Not applicable

## Supplementary information

### Supplementary data

**Supp Fig. 1:**
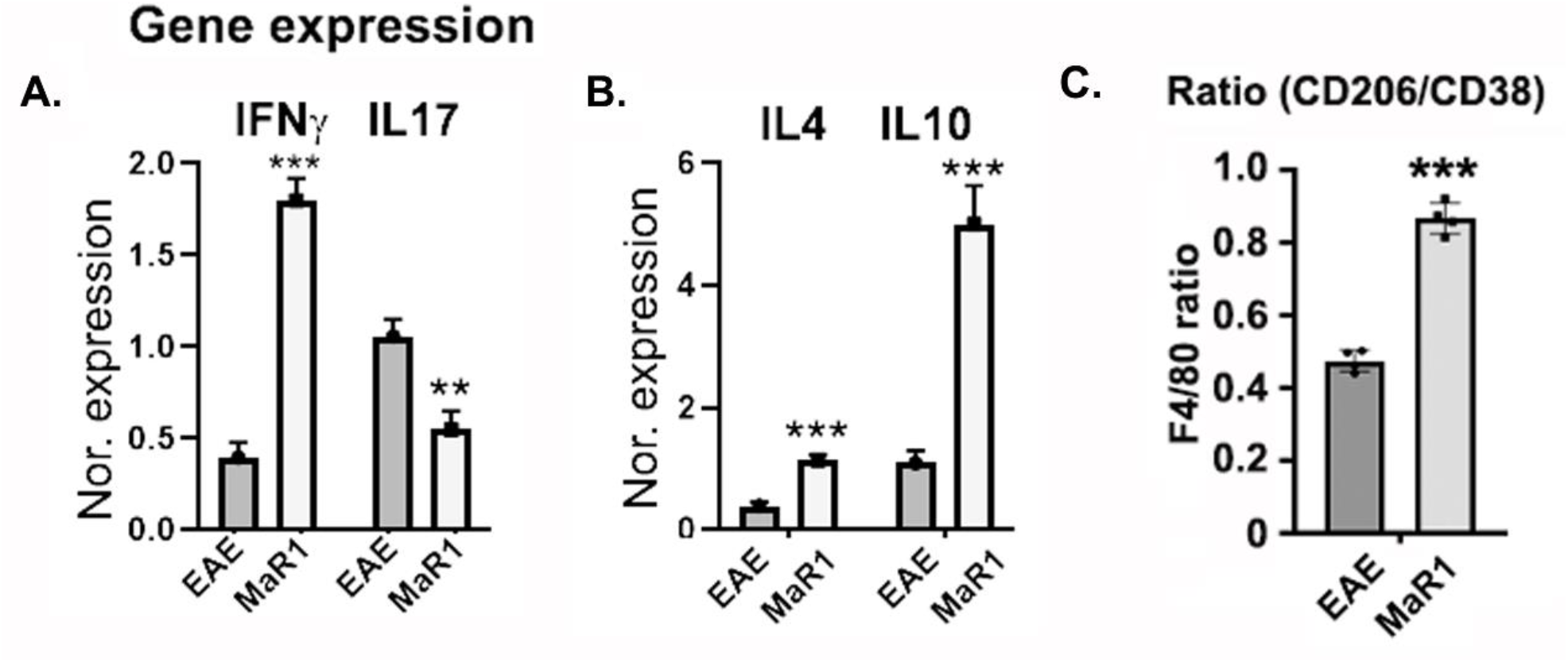
MaR1 modulated the expression of pro- and anti-inflammatory cytokines in spleen/LN cells and altered the phenotype of macrophages in the CNS. **A-B**. RNA was isolated from spleen/LN cells from mice with untreated EAE and MaR1 after 24 h of antigen stimulation, after which the expression of IFN*γ*, IL17a, IL4 and IL10 was examined (n=3). **C**. CNS tissues (brain and spinal cord together) were processed, and CD206+ and CD38+ F4/80+ cells in the CNS of treated and untreated RR-EAE (n=4) were examined. The ratio of CD206/CD38+ macrophages was plotted to determine the macrophage phenotype (n=4). ***, P<0.01.

**Supp Figure 2:**
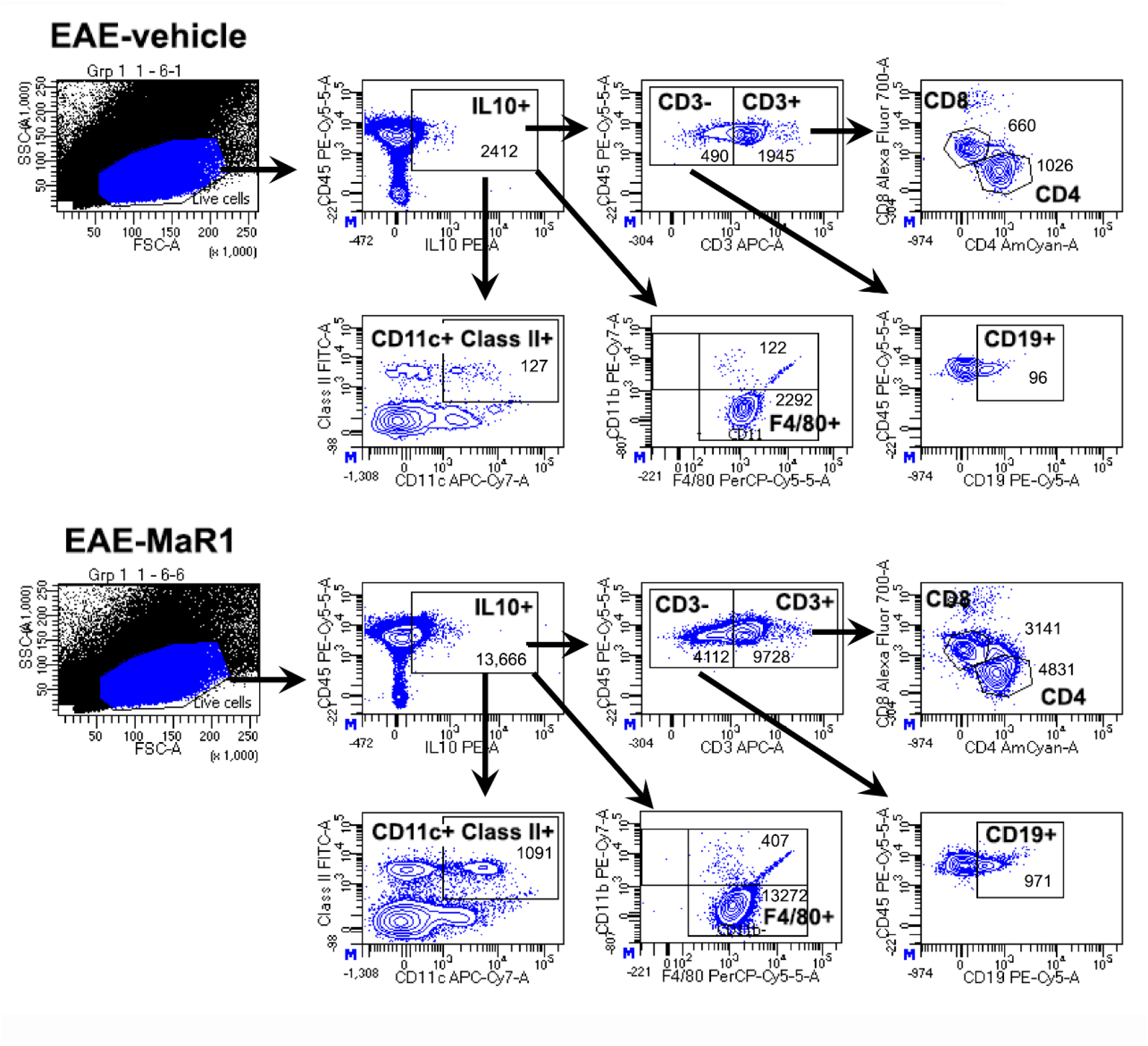
Gating strategy for examining IL10-expressing immune cells. Using a Percoll density gradient, brain and spinal cord single-cell suspensions from EAE and MaR1-treated EAE mice were generated and subjected to flow cytometry analysis. Before identifying IL10-expressing lymphoid cells, CD3+ populations were gated from IL10+ populations. CD4+ and CD8+ T cells were identified in CD3+ populations, while B cells were identified in CD3-populations. To identify IL10-expressing myeloid populations, CD11c+ClassII+ populations were gated for inflammatory dendritic cells, whereas CD11b+ F4/80+ populations were gated for macrophages from IL10+ populations.

**Supp Fig 3:**
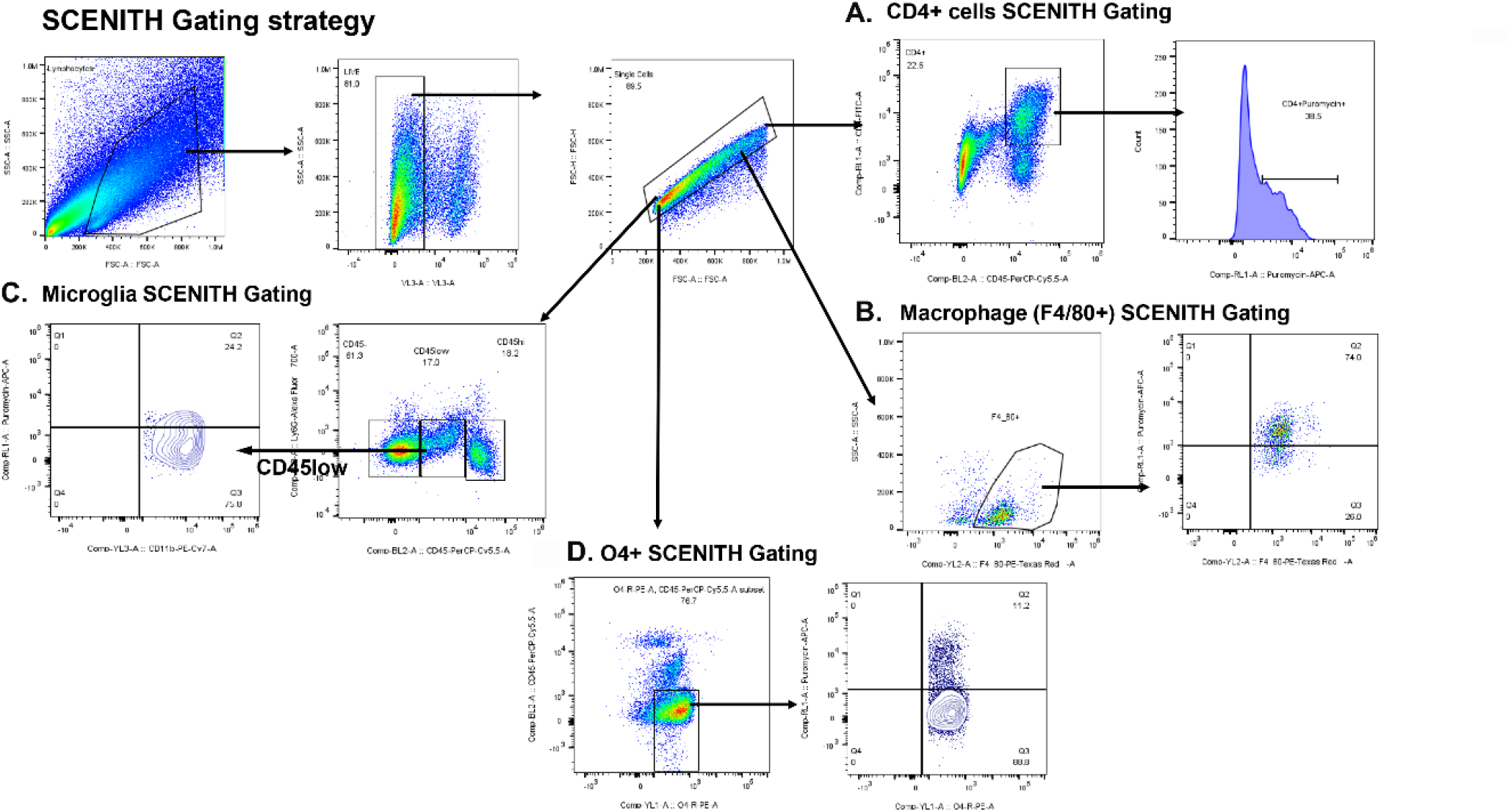
Gating a strategy for immunophenotyping using SCENITH. The brain and spinal cord single-cell suspensions were stained for lymphoid and myeloid markers, as described in the methods section. To eliminate doublets and dead cells, the Live_Dead-ve population was gated, and the FSC-A vs FSC-H data were plotted from negative populations, identifying these populations as live cells, which were used in all subsequent analyses. CD4+ T-cell populations were identified by double gating of CD45+CD4+ cells, and puromycin-positive cells were gated from this population. Microglial populations were gated based on CD45lowCD11b+ and puromycin double-positive cells. To detect puromycin+ macrophage populations, CD45+F4/80+ cells were gated for total macrophages, followed by puromycin-positive populations.

**Supp Fig 4:**
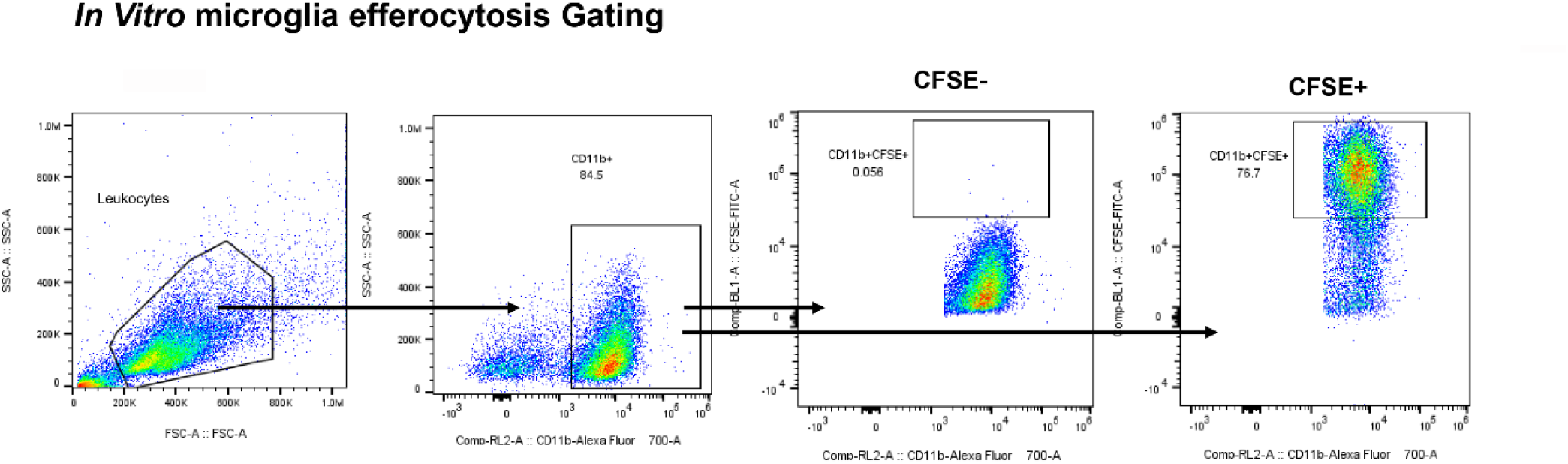
Gating strategy for e*x vivo* efferocytosis analysis. A pseudocolor map was used to quantify total efferocytosis in macrophages and microglia from mice subjected to EAE and treated with MaR1. Double-positive gating of CD11b+CFSE+ populations revealed apoptotic cell engulfment by macrophages and microglia, whereas CD11b+CFSE-populations were identified as nonefferocytotic macrophages and microglia.

